# Temporal Structure in Resting-State fMRI under Marginal and Cross-Spectral Surrogate Controls

**DOI:** 10.64898/2026.11.04.763876

**Authors:** Aleksandr Volkov, Yury Maximov, Tamoghna Chattopadhyay, Paul M. Thompson, Ildar Gabitov

**Affiliations:** PowerMechaniX LLC, Albuquerque, New Mexico, USA; Imaging Genetics Center, Mark & Mary Stevens Neuroimaging and Informatics Institute, Keck School of Medicine, University of Southern California, Marina del Rey, CA, USA; Department of Mathematics, University of Arizona, Tucson, Arizona 85721, USA

**Keywords:** Resting-state fMRI, surrogate data, sample entropy, finite-scale structure, multifractal analysis

## Abstract

Resting-state functional MRI is commonly analyzed using correlations between regional signals and their frequency spectra, which characterize average connectivity and spectral properties but may miss other aspects of temporal organization. Measures of temporal complexity offer complementary information about the recurrence of signal patterns, how fluctuations vary across timescales, and signal roughness. These properties may reveal disease-related alterations in brain activity, offering opportunities for biomarker discovery and insights into mechanisms of neurological and psychiatric disorders. Here we examined sample entropy (SampEn), multifractal de-trended fluctuation analysis (MFDFA), and Higuchi curve shapes in 732 scans from 270 ADNI participants, including cognitively normal individuals, people with mild cognitive impairment, and people with dementia. Using the seven Yeo resting-state networks (Yeo-7), we compared recordings with randomized signals preserving either each network’s amplitude distribution and approximate spectrum (IAAFT), or the complete cross-spectrum across networks. If these properties fully explained temporal complexity, real recordings should resemble their randomized references. We therefore counted how often each real recording was the strongest outlier among itself and 20 references. This occurred in 21.7%, 12.7%, and 44.2% of 5,124 network-visits for SampEn, MFDFA, and Higuchi, versus 5.5%, 5.1%, and 6.3% in surrogate-only controls. Cross-spectrum-preserving references yielded 39.8%, 17.5%, and 49.8%. Simulations showed that changes in sampling or signal processing, including abrupt changes in signal amplitude, can produce similar differences without changing the underlying dynamics. The measures showed repeatability across visits but did not improve retrospective prediction of changes in Clinical Dementia Rating Sum of Boxes (CDR-SB) scores. These findings identify temporal features not reproduced by the tested spectral controls and provide a basis for evaluating their potential as disease biomarkers.

## 1 INTRODUCTION

Resting-state functional MRI is a noninvasive method to study spontaneous brain activity and how its organization changes in neurological and psychiatric disorders [1]. Such changes may provide biomarkers of disease and insights into underlying brain mechanisms. Most analyses summarize activity using correlations between time-series between pairs of brain regions or using their frequency spectra. These measures may miss how fluctuations vary across timescales, and how irregular the signals are. Sample entropy measures the recurrence of short patterns, multifractal detrended fluctuation analysis (MFDFA) characterizes fluctuations across timescales, and Higuchi curves measure signal roughness at different scales. These complementary measures may reveal disease-related information not captured by conventional analyses [2, 3]. However, we first need to establish what additional information they measure and whether it reflects genuine temporal organization. Prior studies have used surrogate signals to investigate fMRI multifractality [4], evaluated multifractal features for attention-deficit/hyperactivity disorder classification [5], and examined the effects of sampling and recording duration [6]. Here, we analyze the full shapes of MFDFA and Higuchi curves alongside sample entropy, rather than reducing each curve to a single fitted slope. This retains information across time scales without assuming a power law. We compare these measures with randomized reference signals that preserve selected statistical and spectral properties, to determine what temporal information remains unexplained [7].

To determine whether temporal complexity measures capture information missed by conventional fMRI analyses, we analyzed 732 ADNI scans using the Yeo-7 resting-state networks (RSNs), seven large-scale groups of brain regions whose activity is correlated at rest [8]. We averaged signals within each network and compared the resulting time series with two complementary types of randomized reference signals. Iterative amplitude-adjusted Fourier transformation (IAAFT) preserves each network’s amplitude distribution and approximately its frequency spectrum [9, 10], allowing us to test for additional temporal structure. Common-phase randomization preserves the complete cross-spectrum, including frequency-dependent relationships between networks, while allowing amplitude distributions to change [11]. Together, these comparisons help determine which temporal features are not explained by conventional spectral properties. Finally, we examined repeatability across visits and prediction of clinical change to evaluate the potential of these measures as disease biomarkers.

## 2 COHORT & TEMPORAL MEASUREMENTS

### 2.1 Participants & Study Design

To examine temporal fMRI features across stages of cognitive impairment, we selected cognitively normal (CN), mild cognitive impairment (MCI), and dementia participants from the Alzheimer’s Disease Neuroimaging Initiative (ADNI) [12]. Of 1,764 participants with 4,347 BOLD fMRI series, 1,698 met clinical and imaging criteria, including diagnosis within 90 days of MRI, and adequate time-series length. Of these, 907 had longitudinal data and 903 met additional requirements for matching.

To minimize demographic and acquisition differences, we matched each of 90 eligible dementia participants to one CN and one MCI participant, requiring the same number of visits and similar demographics. The final cohort comprised 270 participants (90 per group) and 732 fMRI scans. All 732 scans were included in the primary analysis. Averaging regional signals within the seven Yeo resting-state networks yielded 5,124 network-level time series. To account for repeated measures within individuals, bootstrap confidence intervals were calculated by resampling participants.

### 2.2 Network Time Series

We extracted regional BOLD time series using the Schaefer-400 atlas [13], with parcel labels assigned by nearest-neighbor interpolation. Within each parcel, we averaged signals from voxels with valid values throughout the scan, without restricting them to the fMRIPrep brain mask [14]. We removed initial non-steady-state volumes and averaged parcel time series with equal weights to obtain one time series for each of the seven Yeo resting-state networks. To preserve the original temporal properties of the signals, this analysis used data before nuisance regression, without additional filtering, temporal interpolation, volume censoring, or standardization. Each scan retained its original repetition time (TR).

### 2.3 Sample Entropy and Finite Curve Shapes

Let *Y*_*i*_(*t*) denote the mean BOLD signal in Yeo network *i* = 1,… ,7 at fMRI time point *t*. For each network separately, sample entropy measures how regularly short patterns of activity recur [15]. We compare the signal at two different starting times, *t* and *s*. If the signals are similar at both t and s, and again at t+1 and s+1, do they also remain similar at t+2 and s+2? More consistent repetition indicates lower entropy. For sequences of length m = 2 or 3, we define their difference as

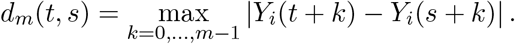

Sequences match when *d*_*m*_(*t, s*) ≤ 0.2*σ*_*i*_, where *σ*_*i*_ is the standard deviation of the network signal. Using starting times 0, … , N − 3, let B count matching pairs of length 2, and A count those that still match at the third time point. The ratio A/B measures how often similar patterns continue to match, giving sample entropy, *S*_*i*_ = log(*A/B*). Self-comparisons were excluded, but overlapping sequences were allowed. Entropy was undefined if either count was zero, and estimates were less precise for short scans [16].

Multifractal de-trended fluctuation analysis (MFDFA) measures how signal fluctuations change across time scales, revealing differences in temporal organization [17]. We integrated each mean-centered network time series and removed linear trends within nonoverlapping segments. The resulting fluctuation measure *F*_*q*_(*s*) depends on segment length s, with q = −2, 0, 2, 4 giving different weights to small and large fluctuations (using a logarithmic definition at q = 0). We retained 12 values relative to a reference scale of eight time points:

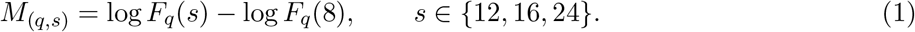

Higuchi’s method, originally developed to estimate the fractal dimension of a time series, measures how its apparent length changes when examined at different temporal resolutions [18, 19]. This measures signal roughness across time scales. Using lag- and offset-normalized curve lengths L(k), we retained 11 values:

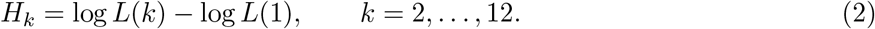

Retaining these curves rather than fitting a single slope preserves differences across timescales without assuming a power law.

## 3 SURROGATE REFERENCES & CALIBRATION

### 3.1 Marginal and Spectral Constraints

To test whether temporal complexity measures capture information beyond conventional spectral properties, we compared fMRI signals with two types of randomized surrogates. We generated 20 iterative amplitude-adjusted Fourier transform (IAAFT) surrogates per network and visit. Starting from shuffled signals, 25 iterations alternately matched the original Fourier amplitudes and signal-value distribution. The surrogates preserved the amplitude distribution exactly and the power spectrum approximately. We recorded the remaining spectral error and repeated the analysis using cases where all 20 errors were less than 10^−3^. Second, to preserve relationships between networks, we generated 20 multichannel surrogates per scan by applying the same random Fourier phase *φ*(f) to all seven networks at each frequency: 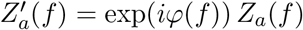, and

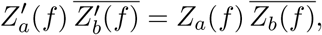

where *Z*_*a*_(*f*) is the Fourier coefficient for network a; the overbar denotes complex conjugation. Zero-frequency and Nyquist constraints ensured real-valued signals. As the shared phase cancels, all pairwise cross-spectra and full-scan functional connectivity remain unchanged, although amplitude distributions may change. These methods test if temporal complexity captures information beyond individual network spectra and their frequency-dependent relationships.

### 3.2 Ranking and Calibration of IAAFT

To test whether real fMRI signals contain temporal features not reproduced by randomization, we compared each original signal with its 20 IAAFT surrogates. We measured how far each signal’s complexity features differed from the other 20, using a standardized distance:

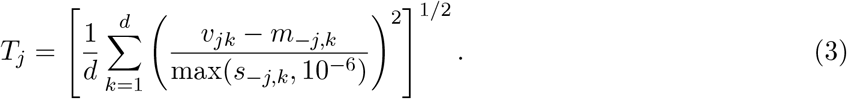

Here, v_j_ is the feature vector for signal j (j = 0 for the original), and m__j,k_ and s__j,k_ are the median and standard deviation of feature k among the other 20 signals. The feature dimensions were d = 1, 12, and 11 for SampEn, MFDFA, and Higuchi. We counted how often the original signal had the highest score, indicating temporal features more different from the reference signals than those signals were from each other.

To check whether randomization could produce extreme scores, we repeated the analysis using an IAAFT surrogate as the original and generating 20 new surrogates from it.

## 4 RESULTS

### 4.1 Observed Ranks and IAAFT Calibration

To test whether real fMRI contains temporal features missed by spectral analysis, we compared each signal with 20 randomized versions. We ranked all 21 signals by how much their complexity measures differed from the others. Here *rank-1* means the real signal was the most different, which should occur in ∼5% of cases if all signals were statistically interchangeable. Across 5,124 network-visits, real signals ranked first in 21.7% of cases for SampEn, 12.7% for MFDFA, and 44.2% for Higuchi (95% confidence intervals: 19.6–24.1%, 11.5–14.2%, and 42.0–46.4%). When the experiment was repeated using randomized signals, these fractions fell to 5.5%, 5.1%, and 6.3% (Table 1, Fig. 2). Thus, real fMRI exhibited temporal features, particularly Higuchi curve shapes, that were not reproduced by spectral randomization.

**Table 1.**
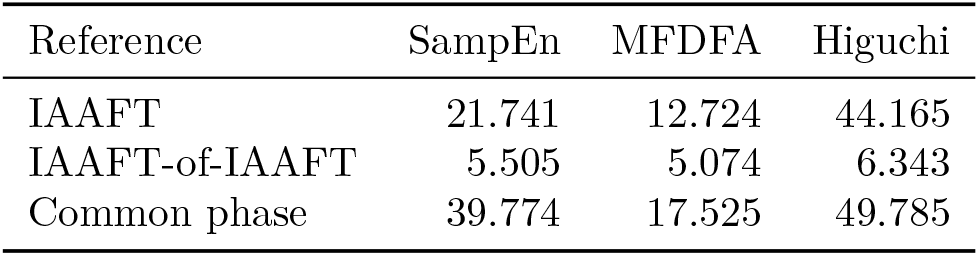
Percentage of signals with the most extreme complexity score among 21 signals, comparing real fMRI with randomized references and a surrogate-only control.

**Figure 1.**
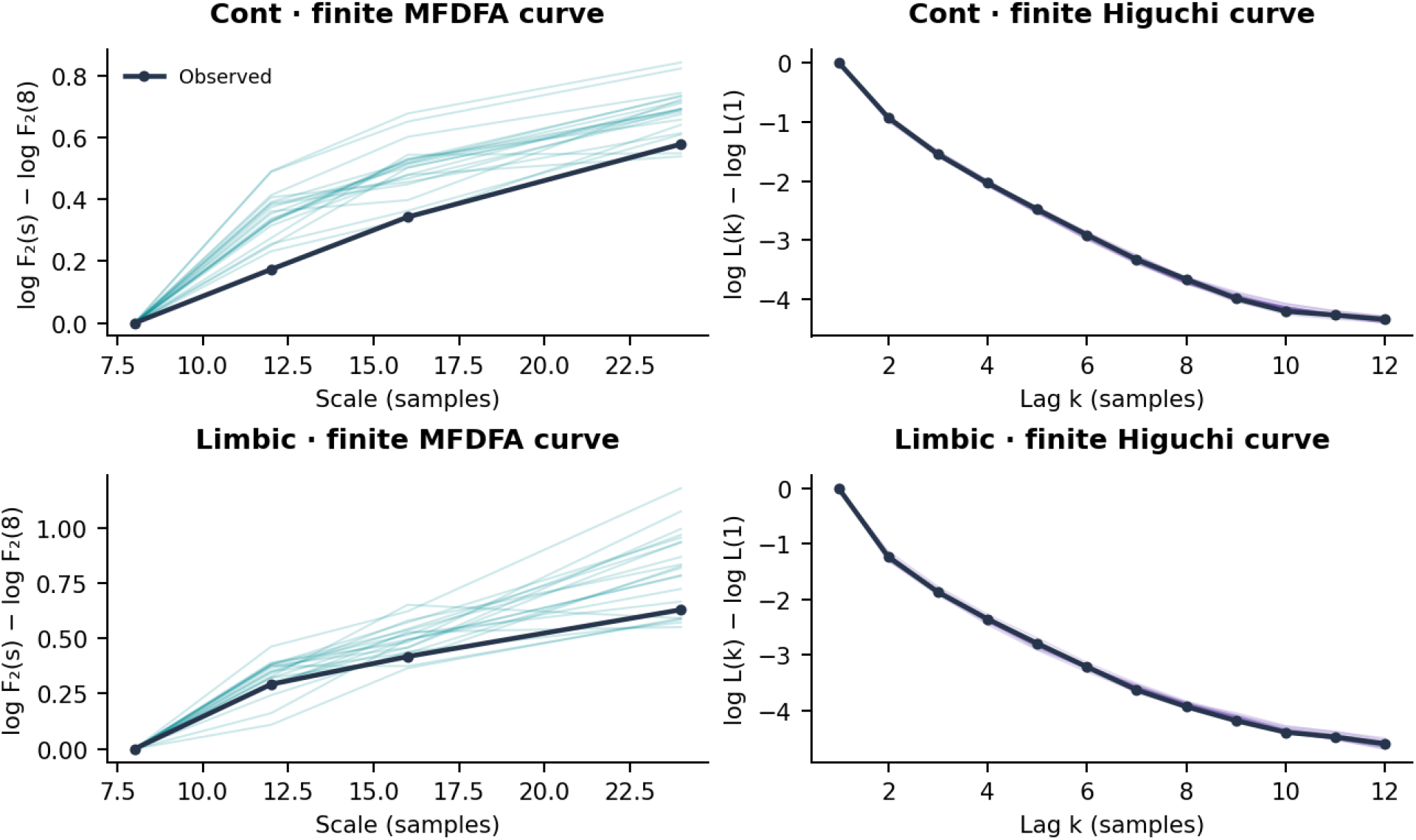
Temporal complexity curves for the Control and Limbic networks in an example fMRI scan. Dark lines show observed data; light lines show 20 IAAFT surrogate signals. MFDFA panels (left) show q = 2; Higuchi panels (right) show the full curves, normalized to zero at lag 1. Timescales are in samples (TR = 3 s).

**Figure 2.**
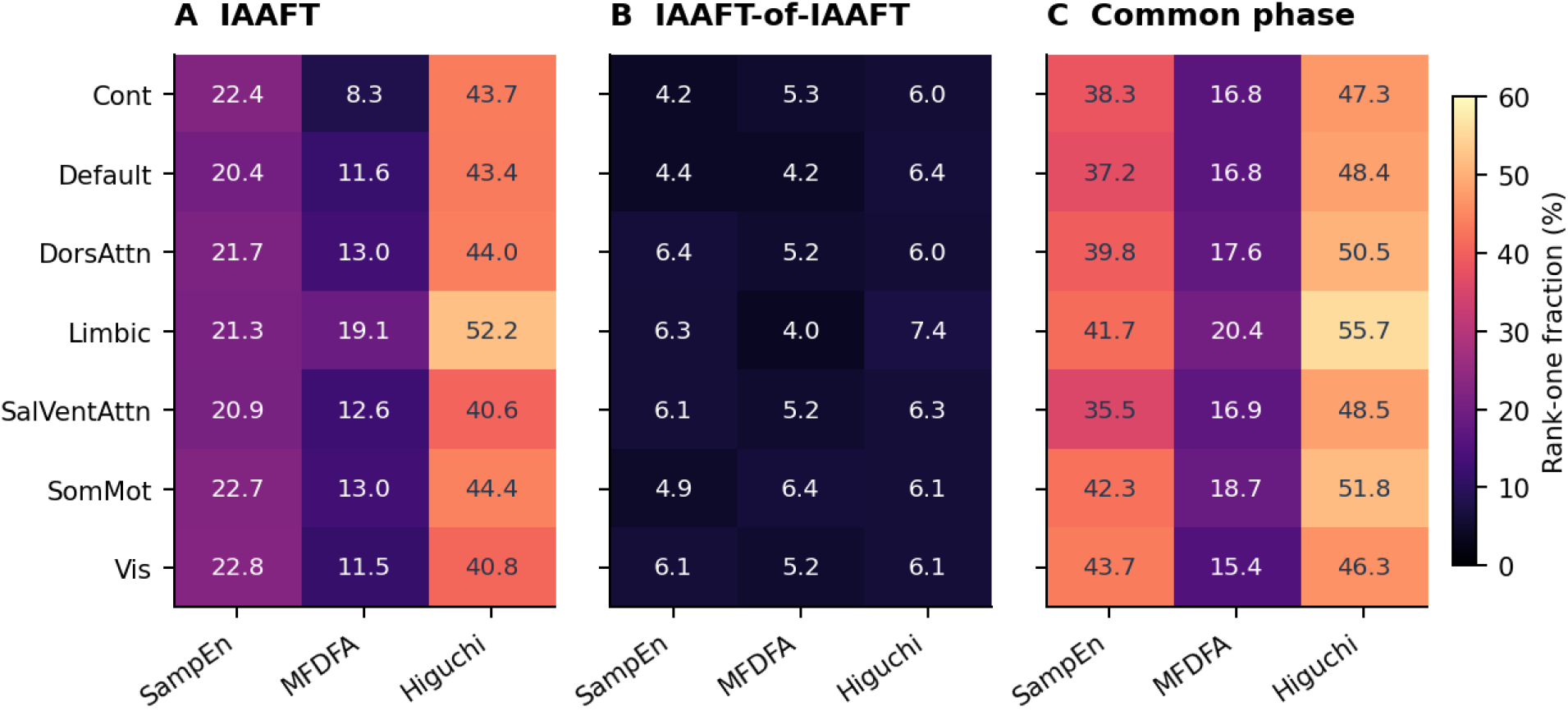
Percentage of fMRI network measures that differed more from the surrogate distribution than any of 20 randomized references, for each resting-state network and complexity measure. Middle panel uses randomized signals instead of real fMRI data to assess if the randomization method produces such differences.

The median IAAFT Fourier-amplitude error was 7.25 × 10−5 (maximum 0.024). The stricter all-reference error criterion retained 4,969 network-visits and fractions of 20.7%, 12.6% and 44.0%, close to the main values. SampEn was below its reference median in roughly 68–76% of visits within each network. The rank measures outlyingness; the curve differences need not all represent increased complexity.

### 4.2 Temporal Differences at a Fixed Cross-Spectrum

Even when the 7-network cross-spectrum was preserved, the original fMRI signals had more extreme complexity scores than all 20 randomized references in 39.8% (SampEn), 17.5% (MFDFA), and 49.8% (Higuchi) of network-visits. All measures were valid; cross-spectrum preservation was nearly exact (maximum relative error 9.78 × 10-16). Thus, the measures detected temporal differences not explained by the preserved spectral relationships. However, randomization changed the signal-value distributions (median RMS difference, 0.165 SD). Results depended on scan length: SampEn fractions were 27.9% for shorter scans (N ≤ 197; 3,115 network-visits) versus 58.1% for longer scans (N > 197; 2,009 network-visits). These groups also differed in acquisition characteristics and participant composition.

### 4.3 Observation Effects at Fixed Latent Dynamics

For temporal complexity measures to be useful as disease biomarkers, they must distinguish changes in brain activity from differences caused by MRI acquisition or signal processing [20, 21]. We therefore tested if identical underlying dynamics could produce different complexity measurements when sampled or processed differently. We generated 32 realizations of stationary Gaussian processes, tested at ten combinations of scan length and repetition time (TR), with four signal transformations, giving 1,280 observations.

The effects were sometimes striking. At N = 480 and TR = 0.607 s, doubling the signal amplitude halfway through the scan increased extreme SampEn rankings from 6.7% to 99.1%. At N = 197 and TR = 3 s, a cubic transformation increased extreme Higuchi rankings from 7.1% to 41.1% (Fig. 3). Other conditions showed little change, while short scans reduced measurement reliability. Thus, apparent differences in temporal complexity can arise without changes in underlying brain dynamics, highlighting an important potential confound in biomarker studies.

**Figure 3.**
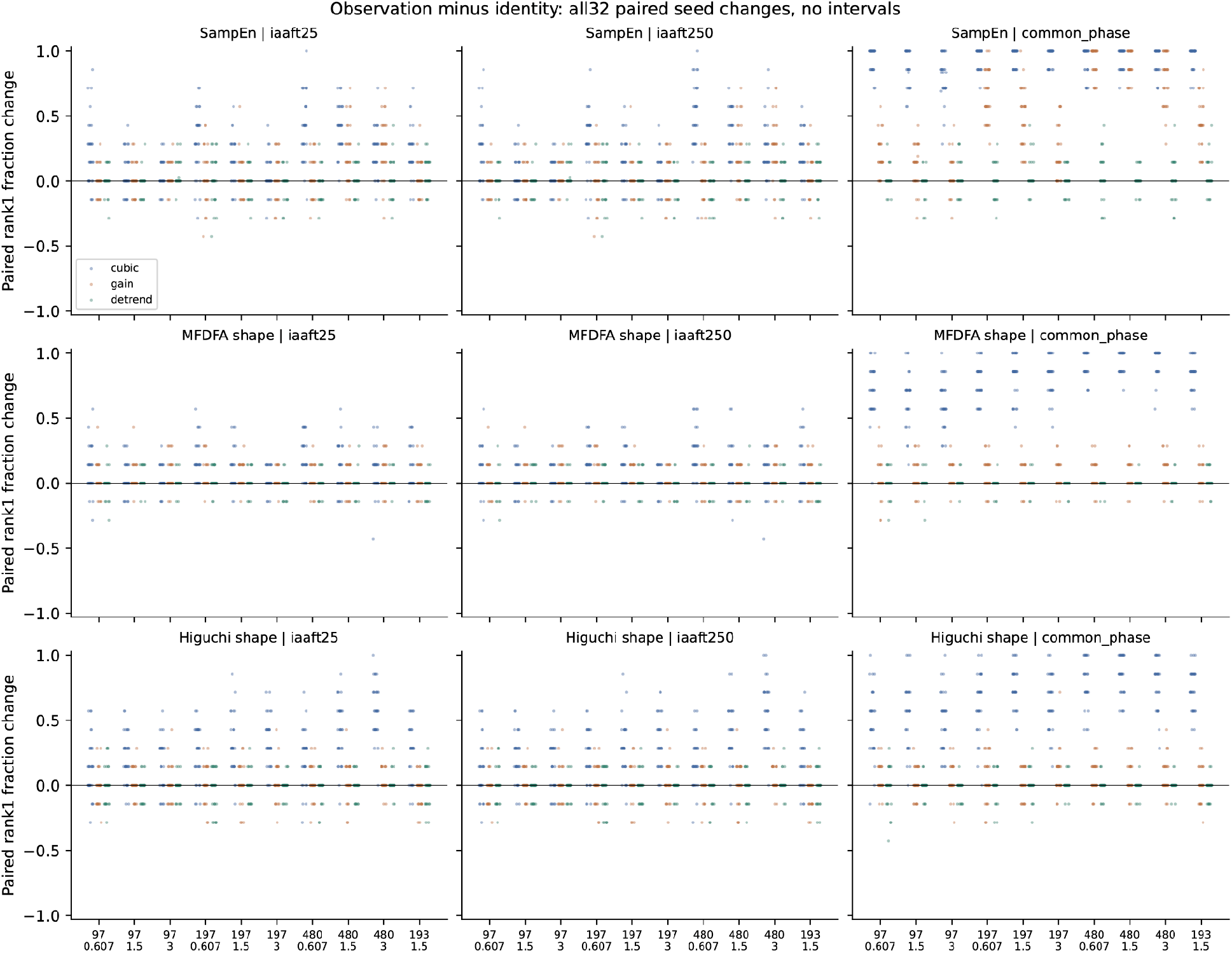
Effects of signal transformations on temporal complexity, relative to unchanged simulated signals. Each point summarizes valid network measurements from one simulation. Colors distinguish cubic transformation, mid-scan amplitude increase, and detrending. Panels compare complexity measures and surrogate methods; positive, negative, and zero values indicate increases, decreases, or no change in extreme rankings.

### 4.4 Individual Correspondence and Clinical Change

For potential biomarker use, temporal complexity measures should show individual consistency and clinical relevance. Across 409 visit pairs from 246 participants with matching TRs, measurements were more similar within individuals than in matched comparisons in 62.6% (SampEn), 60.6% (MFDFA), and 71.9% (Higuchi) of pairs, comparable to conventional measures (65.0–73.1%). We next examined 264 intervals from 190 participants, retrospectively estimating the magnitude of intervisit change in Clinical Dementia Rating Sum of Boxes (CDR-SB) [22]. A model using functional connectivity, spectral features, and signal power achieved a mean absolute error (MAE) of 0.969 points. Adding SampEn, MFDFA, or Higuchi increased MAE by 0.018, 0.066, and 0.030 points, respectively; clinical covariates alone achieved 0.934. Thus, these measures showed individual consistency, but no additional clinical predictive value in this pilot analysis.

## 5 DISCUSSION & CONCLUSION

This pilot study shows that resting-state fMRI contains temporal features not reproduced by conventional spectral controls. Sample entropy and the full MFDFA and Higuchi curves distinguished real fMRI signals from randomized references, even when the complete cross-spectrum across seven resting-state networks was preserved. Extending earlier work on fMRI multifractality [4], we retained information across timescales rather than reducing each curve to a single slope. Combining complementary surrogate methods with a calibration test also enables more rigorous assessment of what these measures capture beyond conventional functional connectivity and spectral analysis.

Some limitations remain. Simulations show that sampling and processing can alter apparent temporal complexity without changing the underlying dynamics. These effects must be considered when interpreting biological differences, as with other dynamic fMRI metrics [20, 23]. The measures were consistent across visits but did not improve prediction of clinical change. However, this pilot study offers a basis for larger studies of fMRI temporal complexity in neurology and psychiatry.

## 6 CODE & DATA AVAILABILITY

Analysis code is available on GitHub: https://github.com/yury-rd/temporal-surrogates

## 7 COMPLIANCE WITH ETHICAL STANDARDS

We analyzed previously collected, publicly available ADNI data, without recruiting participants or acquiring new scans. For this analysis of public, de-identified anonymized data, no additional institutional review board approval was required. The synthetic analyses contained no human data.

## Acknowledgments

Data used in preparation of this article were obtained from the Alzheimer’s Disease Neuroimaging Initiative (ADNI) database (adni.loni.usc.edu). As such, the investigators within the ADNI contributed to the design and implementation of ADNI and/or provided data but did not participate in the analysis or writing of this report. A complete listing of ADNI investigators can be found in the ADNI investigator acknowledgment list.

The ADNI imaging and clinical data came from the Alzheimer’s Disease Neuroimaging Initiative database. ADNI began in 2003 as a public-private partnership led by Michael W. Weiner, MD. It brings together longitudinal imaging, biomarkers, and clinical and neuropsychological assessments to study progression from mild cognitive impairment to Alzheimer’s disease.

The National Institute on Aging funds ADNI data collection and sharing through National Institutes of Health grant U19 AG024904 to the Northern California Institute for Research and Education. ADNI has also received support from the National Institute of Biomedical Imaging and Bioengineering, the Canadian Institutes of Health Research, and private contributors through the Foundation for the National Institutes of Health. These contributors include AbbVie, Alzheimer’s Association, Alzheimer’s Drug Discovery Foundation, Araclon Biotech, BioClinica, Inc., Biogen, Bristol-Myers Squibb Company, CereSpir, Inc., Cogstate, Eisai Inc., Elan Pharmaceuticals, Inc., Eli Lilly and Company, EuroImmun, F. Hoffmann-La Roche Ltd and its affiliated company Genentech, Inc., Fujirebio, GE Healthcare, IXICO Ltd., Janssen Alzheimer Immunotherapy Research & Development, LLC., Johnson & Johnson Pharmaceutical Research & Development LLC., Lumosity, Lundbeck, Merck & Co., Inc., Meso Scale Diagnostics, LLC., NeuroRx Research, Neurotrack Technologies, Novartis Pharmaceuticals Corporation, Pfizer Inc., Piramal Imaging, Servier, Takeda Pharmaceutical Company, and Transition Therapeutics.

## References

[1] M. D. Fox and M. E. Raichle, “Spontaneous Fluctuations in Brain Activity Observed with Functional Magnetic Resonance Imaging,” Nature Reviews Neuroscience, vol. 8, no. 9, pp. 700–711, 2007. doi: 10.1038/nrn2201.

[2] E. Fiorenzato, S. Moaveninejad, L. Weis, R. Biundo, A. Antonini, and C. Porcaro, “Brain Dynamics Complexity as a Signature of Cognitive Decline in Parkinson’s Disease,” Movement Disorders, vol. 39, no. 2, pp. 305–317, 2024. doi: 10.1002/mds.29678.

[3] Z. Wang, for the Alzheimer’s Disease Neuroimaging Initiative, “Brain Entropy Mapping in Healthy Aging and Alzheimer’s Disease,” Frontiers in Aging Neuroscience, vol. 12, article 596122, 2020.

[4] Sihai Guan, Dongyu Wan, Yanmiao Yang, and Bharat Biswal, “Sources of multifractality of the brain rs-fMRI signal,” Chaos, Solitons & Fractals, vol. 160, pp. 112222, 2022. doi: 10.1016/j.chaos.2022.112222.

[5] Mengyunqiu Zhang, Sei-ichiro Kamata, and Pengfeng Lu, “ADHD Classification Method based on Spatio-Temporal Multifractal Features Using Attention-Based Pyramid Feature Mixer Modules,” in 2024 IEEE International Symposium on Biomedical Imaging (ISBI), 2024, pp. 1–5.

[6] Sihai Guan, Chun Meng, and Bharat Biswal, “Fractal complexity of spontaneous brain activity and the effect of scanning parameters,” Advances in Continuous and Discrete Models, vol. 2025, no. 1, pp. 42, 2025. doi: 10.1186/s13662-025-03910-2.

[7] J. Theiler, S. Eubank, A. Longtin, B. Galdrikian, and J. D. Farmer, “Testing for Nonlinearity in Time Series: The Method of Surrogate Data,” Physica D: Nonlinear Phenomena, vol. 58, nos. 1-4, pp. 77–94, 1992.

[8] B. T. T. Yeo, F. M. Krienen, J. Sepulcre, M. R. Sabuncu, D. Lashkari, M. Hollinshead, J. L. Roffman, J. W. Smoller, L. Zöllei, J. R. Polimeni, B. Fischl, H. Liu, and R. L. Buckner, “The Organization of the Human Cerebral Cortex Estimated by Intrinsic Functional Connectivity,” Journal of Neurophysiology, vol. 106, no. 3, pp. 1125–1165, 2011.

[9] Thomas Schreiber and Andreas Schmitz, “Improved Surrogate Data for Nonlinearity Tests,” Physical Review Letters, vol. 77, pp. 635–638, 1996. doi: 10.1103/PhysRevLett.77.635.

[10] T. Schreiber and A. Schmitz, “Surrogate Time Series,” Physica D: Nonlinear Phenomena, vol. 142, nos. 3–4, pp. 346–382, 2000. doi: 10.1016/S0167-2789(00)00043-9.

[11] Dean Prichard and James Theiler, “Generating surrogate data for time series with several simultaneously measured variables,” Physical Review Letters, vol. 73, pp. 951–954, 1994. doi: 10.1103/PhysRevLett.73.951.

[12] Clifford R. Jack, Jr., Matt A. Bernstein, Nick C. Fox, et al., “The Alzheimer’s Disease Neuroimaging Initiative (ADNI): MRI methods,” J. Magnetic Resonance Imaging, vol. 27, no. 4, pp. 685–691, 2008. doi: 10.1002/jmri.21049.

[13] Alexander Schaefer, Ru Kong, Evan M. Gordon, Timothy O. Laumann, Xi-Nian Zuo, Avram J. Holmes, Simon B. Eickhoff, and B. T. Thomas Yeo, “Local-Global Parcellation of the Human Cerebral Cortex from Intrinsic Functional Connectivity MRI,” Cerebral Cortex, vol. 28, no. 9, pp. 3095–3114, 2018. doi: 10.1093/cercor/bhx179.

[14] O. Esteban, C. J. Markiewicz, R. W. Blair, C. A. Moodie, A. I. Isik, A. Erramuzpe, J. D. Kent, M. Goncalves, E. DuPre, M. Snyder, H. Oya, S. S. Ghosh, J. Wright, J. Durnez, R. A. Poldrack, and K. J. Gorgolewski, “fMRIPrep: A Robust Preprocessing Pipeline for Functional MRI,” Nature Methods, vol. 16, no. 1, pp. 111–116, 2019.

[15] Joshua S. Richman and J. Randall Moorman, “Physiological time-series analysis using approximate entropy and sample entropy,” American Journal of Physiology–Heart and Circulatory Physiology, vol. 278, pp. H2039–H2049, 2000.

[16] J. M. Yentes, N. Hunt, K. K. Schmid, J. P. Kaipust, D. McGrath, and N. Stergiou, “The Appropriate Use of Approximate Entropy and Sample Entropy with Short Data Sets,” Annals of Biomedical Engineering, vol. 41, no. 2, pp. 349–365, 2013. doi: 10.1007/s10439-012-0668-3.

[17] Jan W. Kantelhardt, Stephan A. Zschiegner, Eva Koscielny-Bunde, Armin Bunde, Shlomo Havlin, and H. Eugene Stanley, “Multifractal detrended fluctuation analysis of nonstationary time series,” Physica A, vol. 316, pp. 87–114, 2002. doi: 10.1016/S0378-4371(02)01383-3.

[18] Tomoyuki Higuchi, “Approach to an irregular time series on the basis of the fractal theory,” Physica D, vol. 31, pp. 277–283, 1988. doi: 10.1016/0167-2789(88)90081-4.

[19] L. Pini, S. Moaveninejad, M. Corbetta, and C. Porcaro, “Fractal Dynamics of the Healthy Human Brain: A Subcortical-to-Cortical Gradient of Temporal Complexity,” iScience, vol. 29, no. 6, article 115858, 2026. doi: 10.1016/j.isci.2026.115858.

[20] D. Rubin, T. Fekete, and L. R. Mujica-Parodi, “Optimizing Complexity Measures for fMRI Data: Algorithm, Artifact, and Sensitivity,” PLOS ONE, vol. 8, no. 5, article e63448, 2013. doi: 10.1371/journal.pone.0063448.

[21] J. D. Power, K. A. Barnes, A. Z. Snyder, B. L. Schlaggar, and S. E. Petersen, “Spurious but Systematic Correlations in Functional Connectivity MRI Networks Arise from Subject Motion,” NeuroImage, vol. 59, no. 3, pp. 2142–2154, 2012. doi: 10.1016/j.neuroimage.2011.10.018.

[22] S. E. O’Bryant, S. C. Waring, C. M. Cullum, J. Hall, L. Lacritz, P. J. Massman, P. J. Lupo, J. S. Reisch, and R. Doody, for the Texas Alzheimer’s Research Consortium, “Staging Dementia Using Clinical Dementia Rating Scale Sum of Boxes Scores: A Texas Alzheimer’s Research Consortium Study,” Archives of Neurology, vol. 65, no. 8, pp. 1091–1095, 2008. doi: 10.1001/archneur.65.8.1091.

[23] R. Hindriks, M. H. Adhikari, Y. Murayama, M. Ganzetti, D. Mantini, N. K. Logothetis, and G. Deco, “Can sliding-window correlations reveal dynamic functional connectivity in resting-state fMRI?,” NeuroImage, vol. 127, pp. 242–256, 2016. doi: 10.1016/j.neuroimage.2015.11.055.

